# Striatal enkephalin supports maintenance of conditioned cocaine reward during extinction

**DOI:** 10.1101/2023.02.23.529807

**Authors:** Kanako Matsumura, In Bae Choi, Meera Asokan, Nathan N. Le, Luis Natividad, Lauren K. Dobbs

## Abstract

Drug predictive cues and contexts exert powerful control over behavior and can incite drug seeking and taking. This association and the behavioral output are encoded within striatal circuits, and regulation of these circuits by G-protein coupled receptors affects cocaine-related behaviors. Here, we investigated how opioid peptides and G-protein coupled opioid receptors expressed in striatal medium spiny neurons (MSNs) regulate conditioned cocaine seeking. Augmenting levels of the opioid peptide enkephalin in the striatum facilitates acquisition of cocaine conditioned place preference (CPP). In contrast, opioid receptor antagonists attenuate cocaine CPP and facilitate extinction of alcohol CPP. However, whether striatal enkephalin is necessary for acquisition of cocaine CPP and maintenance during extinction remains unknown. We generated mice with a targeted deletion of enkephalin from dopamine D2-receptor expressing MSNs (D2-PenkKO) and tested them for cocaine CPP. Low striatal enkephalin levels did not attenuate acquisition or expression of CPP; however, D2-PenkKOs showed faster extinction of cocaine CPP. Single administration of the non-selective opioid receptor antagonist naloxone prior to preference testing blocked expression of CPP selectively in females, but equally between genotypes. Repeated administration of naloxone during extinction did not facilitate extinction of cocaine CPP for either genotype, but rather prevented extinction in D2-PenkKO mice. We conclude that while striatal enkephalin is not necessary for acquisition of cocaine reward, it maintains the learned association between cocaine and its predictive cues during extinction learning. Further, sex and pre-existing low striatal enkephalin levels may be important considerations for use of naloxone in treating cocaine use disorder.

## Introduction

Cocaine induces rewarding affective states and drug seeking in part by blocking the dopamine transporter and elevating extracellular dopamine levels and signaling in the striatum. However, the mechanisms that sustain cocaine seeking behavior are complex and involve several non-dopaminergic systems in addition to cocaine-enhanced striatal dopamine transmission. For example, striatal µ opioid receptors (MOR) and the opioid peptide enkephalin have been identified as mechanisms that facilitate cocaine reward, reinforcement, and locomotor sensitization to repeated cocaine^1–4^. Enkephalins, which include met- and leu-enkephalin, are derived from the proenkephalin gene (*Penk*) and have high affinity for MOR as well as δ opioid receptors (DOR). *Penk* is highly expressed in the striatum and localized to dopamine D2 receptor-expressing medium spiny neurons (D2-MSNs). Evidence suggests that D2-MSNs release enkephalin within the striatum, where it suppresses local GABA transmission onto neighboring MSNs by acting via *Gi*-coupled MORs^1^. Shifting the balance of striatal output by dysregulating local striatal GABA transmission is associated with greater locomotor sensitization to repeated cocaine and faster acquisition of cocaine place preference^5^.

Consistent with this mechanism, we found that increasing met-enkephalin levels in the ventral striatum during cocaine place conditioning facilitates acquisition of cocaine place preference^1^. Additionally, long-term cocaine treatment also increases striatal met-enkephalin levels^1^. This raises the interesting possibility that enhanced striatal enkephalin and resulting shifts in striatal output via activation of MORs are important contributors to drug-cue associative learning that underlie conditioned cocaine reward and seeking.

While these data implicate a supportive role for striatal enkephalin in conditioned cocaine seeking, additional studies using pharmacological and genetic approaches suggest enkephalin and MORs are necessary for some aspects of acquisition and expression of cocaine seeking. Systemic administration of the non-selective opioid receptor antagonists naloxone and naltrexone attenuates acquisition and expression of conditioned place preference to cocaine^6–8^. This effect appears to be mediated by MORs, as intracerebroventricular injection of the MOR antagonist CTAP also blocks acquisition of cocaine place preference^9^. Moreover, MORs expressed in specific ventral striatum subregions appear to mediate distinct aspects of cocaine reward. Blocking MORs in the nucleus accumbens core attenuates acquisition, while blocking MORs in the nucleus accumbens shell attenuates expression of cocaine place preference^1,2^. Data from constitutive *Penk* knockout mice suggest enkephalin may be the opioid peptide mediating the MOR response, as these mice have decreased cocaine self-administration, motivational breakpoint for cocaine, and cocaine-induced locomotor sensitization^3,4^. However, enkephalins and MORs are expressed ubiquitously throughout the brain. Thus, it is still unclear whether enkephalin signaling through MORs in the striatum in particular is necessary for these behavioral effects of cocaine. Additionally, there are reported sex differences in humans and rodent models regarding sensitivity to cocaine reward and place conditioning^10^. Female mice acquire and extinguish cocaine place preference faster than male mice^11^, and women are more sensitive to cocaine reward and more susceptible to craving and relapse in response to cocaine-associated cues and contexts than men^10,12^. Despite these reports, the majority of studies examining the role opioid peptides and receptors in cocaine reward and reinforcement did not consider sex as a biological variable in their analyses. Understanding the sex-dependent contribution of the opioid system to conditioned cocaine reward is important from an addiction-treatment perspective, especially considering that opioid pharmacology was recently investigated as a potential therapeutic for cocaine use disorder^13^. The goal of the current study was to determine how enkephalin selectively expressed in the striatum contributes to acquisition, expression, and extinction of cocaine place preference and to investigate potential sex differences in these outcomes.

## Materials and Methods

### Animals

All animal procedures were performed in accordance with guidelines from the University of Texas Austin Institutional Animal Care and Use Committee. Male and female adult mice were used for all experiments. Mice with a homozygous deletion of proenkephalin from striatal D2-MSNs (D2-PenkKO) were generated by crossing *Adora2a-Cre*^*+/-*^ and *Penk*^*f/f*^ mice^14^. We previously used *Adora2a-Cre* line to selectively target D2-MSNs^5^. *Adora2a-Cre*^*-/-*^*;Penk*^*f/f*^ littermates were used as controls. Mice were group-housed in a temperature- and humidity-controlled environment under 12:12hr light/dark cycle (lights on at 06:30) with food and water available *ad libitum*. Behavioral experiments were performed during the light cycle.

### Drugs

Cocaine HCl and naloxone HCl were dissolved in sterile saline and administered intraperitoneally (IP) at a dose of 15 mg/kg and 10 mg/kg, respectively.

### Conditioned Place Preference

Experiments were conducted in a two-chamber apparatus with each chamber having a distinct tactile floor (Grid or Hole). Mice received a pre-conditioning preference test (30 min, saline-primed), 8 conditioning days (15 min), and 3 post-conditioning preference tests (30 min). During conditioning, mice received IP cocaine (15 mg/kg) or saline on alternating days and were confined to one chamber. Drug-floor pairing, drug side, and treatment order were counterbalanced across animals. For preference tests, mice received saline (pretest, post-test 1-2) or a 20-min naloxone pretreatment (10 mg/kg; post-test 3) and were allowed to freely explore both chambers. After post-test 3, one cohort received 8 forced extinction days (15 min) where mice received saline prior to placement on their previous cocaine (conditioned stimulus +, CS+) or saline (conditioned stimulus -, CS-) floor. To test whether blocking opioid receptors facilitates extinction, one cohort received naloxone 20 min before placement on the CS+ floor. Extinction was measured in 2, saline-primed preference tests (30 min) after 4 and 8 extinction days. Preference was determined by calculating the percent time spent on the cocaine-paired floor and the “Time on Grid”:

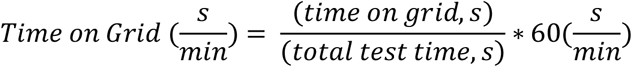

A significant difference in the “Time on Grid” between the conditioning subgroups (Grid + cocaine and Hole + cocaine) indicates preference. For instance, mice that received cocaine on the grid floor are expected to have higher “Time on Grid” than mice that received cocaine on the hole floor^15^. This provided an additional measure of preference and was used when subgroups had sufficient power to detect differences (Pre-Test – Post Test 3). Additionally, to control for baseline differences in preference at Post Test 3, we calculated a preference change score by subtracting the percent time on during the extinction tests from the percent time on Post Test 3.

### Quantitative Polymerase Chain Reaction

D2-PenkKO and *Penk*^*f/f*^ mice were deeply anesthetized with isoflurane, decapitated, and brains dissected on ice. Tissue was homogenized, RNA was extracted, and cDNA was synthesized. Relative mRNA expression of proenkephalin, µ opioid receptor, δ opioid receptor, prodynorphin, and β-actin were determined with TaqMan Gene Expression Assays using a CFX384 Real-Time System (initial hold at 95°C for 20 sec, 40 cycles at 95°C for 1 sec and 60°C for 20 sec). Samples were run in triplicate, and negative controls were run in parallel. Relative expression was calculated using the ^∆∆^Ct method.

### RNAscope Fluorescent Multiplex Assay

D2-PenkKO and *Penk*^*f/f*^ mice were deeply anesthetized with isoflurane, decapitated, and brains were removed and flash-frozen in isopentane on dry ice. Brains were sectioned (12 µm) on a cryostat and mounted on slides. Sections were fixed in 4% PFA in PBS (15 min; 4°C), washed in PBS (2 × 1 min; RT), and dehydrated in ethanol (50%, 1 × 5min; 70%, 1 × 5 min; 100%, 2 × 5 min; RT). Slices were incubated in 100% ethanol at -20°C overnight. After drying (5 min, RT), sections were treated with Protease Pretreat-4 (ACD Bio) and incubated at RT (30 min). Sections were then washed (ACD Bio wash buffer, 2 × 1 min) and incubated with probes for *Penk* and *Drd2* (40°C; 2 hr), washed again (2 × 2 min), and incubated with amplification buffers for each probe (40°C; 30 min each) with washes in between each amplification buffer (2 × 2 min). Sections were cover slipped (DAPI Fluoromount, SouthernBiotech). Images were acquired with a fluorescent microscope (Zeiss) using a 40x oil immersion objective and analyzed using CellProfiler. The number of *Penk* positive cells and *Penk* puncta within each *Penk* and/or *Drd2* positive cell in the ventral striatum and cortex were counted. A total eight images were taken from three different slices (2-3 images per slice) and averaged within each region of each genotype.

### Immunohistochemistry

D2-PenkKO and *Penk*^*f/f*^ were transcardially perfused with 4% PFA in PBS under isoflurane anesthesia. Brains were fixed overnight in 4% PFA, cryoprotected in 30% sucrose (48 h), and sectioned (30 µm) on a cryostat. Free-floating sections were washed (PBS, 3 × 15 min) and blocked (0.2% Triton-X, 10% normal goat serum in PBS; RT; 1 hr). Sections were incubated with primary antibodies (0.2% Triton-X, 10% normal goat serum in PBS; 4°C; overnight) against metenkephalin (1:100 rabbit anti-met-enkephalin) and Cre-recombinase (1:500 guinea pig anti-cre-recombinase). Slices were washed (PBS, 3 × 15 min) and incubated with secondary antibodies (0.2% Triton-X, 1% normal goat serum, PBS; RT; 2hr) conjugated with AlexaFluor-568 (1:1000 goat anti-rabbit) and AlexaFluor-647 (1:1000 goat anti-guinea pig). Sections were washed (PBS, 3 × 15 min), mounted on slides, and cover slipped (DAPI Fluoromount). Images were acquired on a confocal microscope using a 40x oil immersion objective and processed using ImageJ. Cells were counted for each image and averaged across 9 images (3 images per slice, 3 slices) within each brain region for each genotype.

### Mass spectrometry

Mice (n=9 per genotype) were anesthetized with isoflurane, brains were extracted, and the striatum dissected. Dissected striata were pooled (n=3 per genotype), and samples were prepared on ice in a lysis solution (90% methanol and 0.25% glacial acetic acid) spiked with deuterated leu- and met-enkephalin (YGG-d5F-L, YGG-d5F-M). After samples were homogenized, vortexed, and centrifuged, the supernatant was extracted, vacuum dried, and resuspended (60% acidified methanol in 0.25% glacial acetic acid). Samples were centrifuged on a polyethersulfone 10-kDa molecular weight cut off filter, vacuum dried and resuspended (0.1% formic acid) for mass spectrometry analysis. Pellets were reconstituted (1:100) to estimate total protein. Enkephalins were separated by reverse-phase liquid chromatography (300 µL/min at 45°C) and eluted on a column in a stepwise gradient. DUIS-ESI source was used for ionization with the nebulizing gas flow rate set at 3 L/min. Data acquisition was performed using multiple-reaction monitoring in positive-ion mode (Shimadzu 8060 triple quadrupole mass spectrometer). Based on the abundance in product ion scan, transitions were selected for each native peptide and internal standard and confirmed with in silico MS/MS fragmentation tools. Synthetic peptides were prepared in dilutions (0 to 10 ng/uL) similar to sample preparation and spiked with deuterated internal standard for standard curve generation. Ratio of native to deuterated peptide was plotted on standard curve to generate a fit by linear regression and estimate enkephalin concentrations. Average enkephalin concentration was normalized to total protein.

### Statistics

Place preference data were analyzed using 3-way and 2-way ANOVAs with repeated measures as appropriate (Prism, GraphPad). Multifactorial data sets with missing values were analyzed using a Mixed Effects Model. Gene and protein expression data were analyzed using 2-way ANOVA or unpaired t-tests. Significant interactions were followed up with Sidak-corrected t-test comparisons. Results were considered significant at an alpha of 0.05. All data are presented as mean ± SEM.

## Results

### Validation of cell-specific deletion of enkephalin in D2Penk KO

To examine the role of striatal enkephalin in maladaptive behaviors induced by cocaine, we generated a novel mouse strain with a deletion of the proenkephalin gene (*Penk*) specifically from striatal D2-MSNs (D2-PenkKO). To validate the extent and selectivity of the *Penk* deletion, we first measured *Penk* mRNA expression in the ventral and dorsal striatum (VS, DS) of drug naïve D2-PenkKOs and *Penk*^*f/f*^ using qPCR. *Penk* mRNA was significantly reduced in D2-PenkKO mice compared to *Penk*^*f/f*^ littermate controls (Genotype: F_1, 39_ = 306.8, *p* < 0.0001; VS: t_43_ = 13.32, *p* < 0.0001; DS: t_39_ = 11.62, *p* < 0.0001; Fig. 1A). There was no change in mRNA levels of other common opioid peptides and receptors in the striatum including prodynorphin (*Pdyn*), the MOR or DOR (Fig. 1B-D).

**Figure 1.**
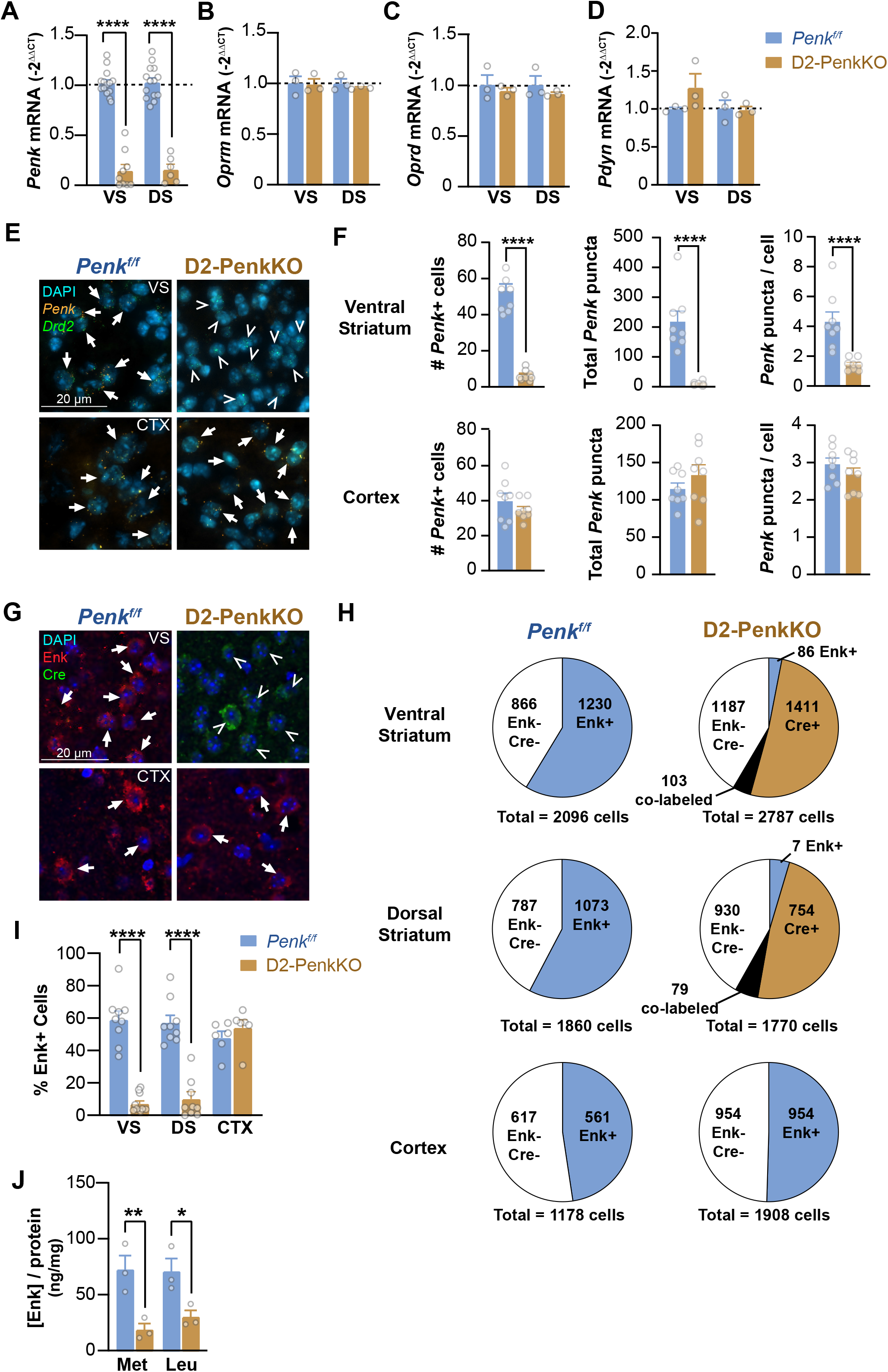
Validation of preproenkephalin gene knockout from D2 receptor expressing striatal medium spiny neurons. (A-D) Significant reduction of preproenekphalin gene (*Penk*: n = 6 – 14 / genotype), but no change in μ opioid receptor (*Oprm*: n = 3 / genotype), δ opioid receptor (*Oprd*: n = 3 / genotype), and preprodynorphin gene (*Pdyn*: n = 3 / genotype) in the ventral (VS) and dorsal (DS) striatum of D2-PenkKO, compared to *Penk*^*f/f*^. (E) Representative images of *in situ* hybridization of *Penk* mRNA (orange), dopamine D2 receptor mRNA (*Drd2*, green), and DAPI (cyan) in the VS (top) and cortex (CTX, bottom) in *Penk*^*f/f*^ (left) and D2-PenkKO (right). Arrows indicate colocalization of DAPI and *Penk*. Triangles indicate colocalization of DAPI and *Drd2*. (F) Significant reduction in total number of *Penk* positive cells (left), total number of *Penk* puncta (middle), and average number of *Penk* puncta per cell (right) in the VS (top), but not the CTX (bottom) of D2-PenkKO compared to *Penk*^*f/f*^ controls (n = 8 / genotype). (G) Representative immunohistochemistry images of VS and CTX in *Penk*^*f/f*^ and D2-PenkKO showing met-enkephalin (Enk, red), Cre recombinase (Cre, green), and DAPI (blue). Arrows indicate colocalization of DAPI and Enk. Triangles indicate colocalization of DAPI and Cre. (H) Proportions of ENK+ (blue), Cre+ (beige), co-labeled (black), and ENK-Cre-cells (white) in the VS (top), DS (middle), and CTX (bottom) of *Penk*^*f/f*^ (left) and D2-PenkKO (right). (I) D2-PenkKO show fewer percent of Enk positive cells in the VS and DS, but not the CTX, compared to *Penk*^*f/f*^ (n = 6 – 9 / genotype). (J) Mass spectrometry showed D2-PenkKO have less met-(Met) and leu-enkephalin (Leu) in the striatum compared to *Penk*^*f/f*^ (n = 3 / genotype). * *p* < 0.05; ** *p* < 0.01; **** *p* < 0.0001 D2-PenkKO vs. *Penk*^*f/f*^

RNAscope was performed to determine whether the deletion *Penk* mRNA was selective for striatal D2-MSNs by assessing the co-expression of *Penk* with the D2 receptor (*Drd2;* Fig. 1E). Relative to *Penk*^*f/f*^ controls, D2-PenkKO mice showed significant reduction in *Penk-Drd2* co-expressing cells (t_12_ = 14.60, *p* < 0.0001), the total number of *Penk* puncta (t_14_ = 5.87, *p* < 0.0001) and the average *Penk* puncta per *Drd2* positive cell (t_14_ = 4.24, *p* < 0.001; Fig. 1F). Importantly, there was no reduction in *Penk* expression within the cortex of D2-PenkKO mice compared to littermate controls, confirming the selectively of the *Penk* knockout for striatal D2-MSNs.

Immunohistochemistry was used to confirm met-enkephalin peptide was significantly reduced in striatal cells. Met-enkephalin positive cells constituted 59.1 ± 5.2% and 57.2 ± 4.6% of all cells in the VS and DS of *Penk*^*f/f*^ controls (Fig. 1I), which is consistent with it selectively being expressed in D2-MSNs^16,17^. In contrast, D2-PenkKO mice had significantly fewer met-enkephalin positive cells in the VS (7.1 ± 1.7%) and DS (10.1 ± 4.4%) (Genotype x Region: F_2, 43_ = 25.41, *p* < 0.0001; DS: t_43_ = 8.18, *p* < 0.0001; VS: t_43_ = 9.76, *p* < 0.0001; Fig. 1I). No reduction in met-enkephalin positive cells was observed in the cortex of D2-PenkKOs. Mass spectrometry of striatal tissue further confirmed that met-enkephalin and leu-enkephalin peptide levels were significantly reduced in D2-PenkKOs compared to *Penk*^*f/f*^ controls (Genotype: F_1, 8_ = 25.41 *p* < 0.01; Fig. 1J).

Since some enkephalin-positive cells remained in the D2-PenkKO striatum, we evaluated the efficiency of recombination by counting Cre and met-enkephalin co-labeled cells. Only a small percentage of Cre-expressing cells were met-enkephalin positive in D2-PenkKOs (VS: 4.2 ± 1.5%, DS: 5.4 ± 2.6%), and Cre was not observed in *Penk*^*f/f*^ controls or in the cortex of D2-PenkKOs (Fig. 1H). Thus, we estimate an ∼95% recombination efficiency in the D2-PenkKO. Taken together, these data provide convergent evidence for a cell-specific deletion of *Penk* from D2-MSNs throughout the entire striatum in D2-PenkKO mice.

### Striatal enkephalin is not required for acquisition or expression of cocaine place preference

We tested whether enkephalin released from D2-MSNs is necessary for acquisition and expression of cocaine CPP. Male and female D2-PenkKO and *Penk*^*f/f*^ mice were conditioned over eight days with preference tests before conditioning (Pretest) and after conditioning trials (Post Test 1 and Post Test 2; Fig. 2A). While deletion of *Penk* had no effect on locomotion during conditioning, cocaine increased locomotion relative to saline trials throughout conditioning (Day x Drug, F_2.164, 131.3_ = 20.63, *p* < 0.0001; SAL vs. COC: t’s = 9.81 – 14.45, *p’s* < 0.0001). Females were also more active than males throughout conditioning for *Penk*^*f/f*^ (Sex: F_1, 41_ = 7.037, *p* < 0.05) and D2-PenkKO (Sex: F_1, 41_ = 17.44, *p* < 0.001; S1A).

**Figure 2.**
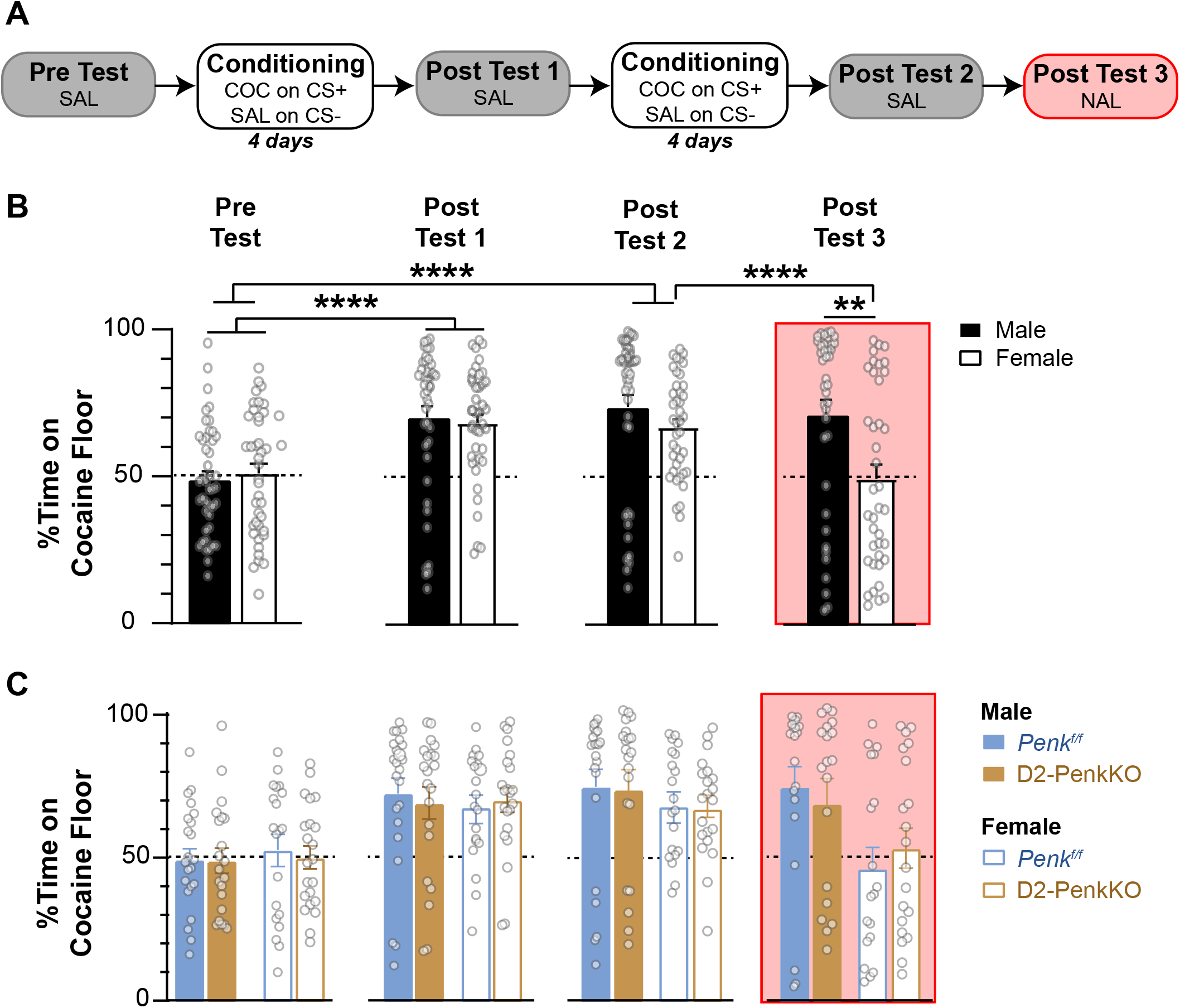
Non-selective opioid antagonist naloxone blocks the expression of cocaine preference selectively in female mice. (A) Experimental timeline of cocaine CPP. Mice received cocaine (15mg/kg; COC) paired with a distinct floor (conditioned stimulus positive, CS+) and saline (SAL) on a different floor (CS-). (B) Both males and females, regardless of genotype, increased the percent time on the cocaine floor after 4 and 8 conditioning days compared to Pre-Test (\*\*\*\**p* < 0.0001; n = 20-23 / genotype). Naloxone pretreatment (10 mg/kg; NAL) blocked expression of cocaine preference in females (n = 44) compared to males (n = 42) at Post Test 3 (** *p* < 0.01) and females at Post Test 2 (**** *p* < 0.0001). (C) Percent time on the cocaine floor is shown for each genotype (*Penk*^*f/f*^ and D2-PenkKO) and sex during Pretest, Post Test 1, Post Test 2, and Post Test 3.

After four conditioning trials, all mice had acquired cocaine CPP, as indicated by a significant increase in the percent time spent on the cocaine-paired floor at Post Test 1 relative to Pretest (Test: F_2, 162_ = 49.66, *p* < 0.0001; t_162_ = 8.39, *p* < 0.0001, Fig. 2B). Preference was maintained for all groups after four additional conditioning trials (t_162_ = 8.84, *p* < 0.0001). These findings were supported by further analysis of the “Time on Grid” (Table 1.) and indicate that neither targeted deletion of *Penk* from striatal D2-MSNs nor sex affects acquisition or expression of cocaine preference.

**Table 1:**
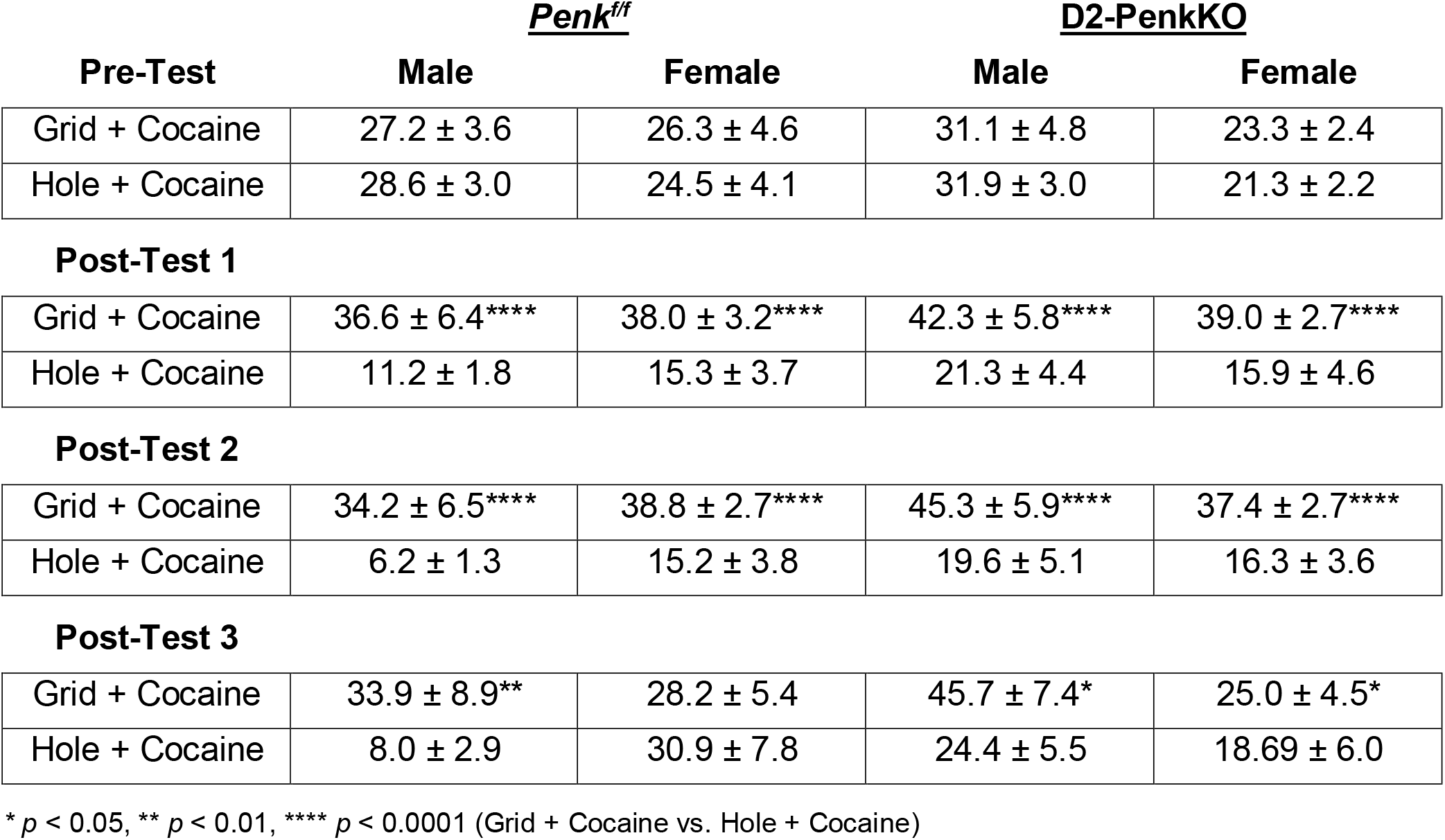
Average time spent on the grid floor during preference tests.

### Acute naloxone blocks expression of cocaine preference selectively in females

Opioid receptor antagonists attenuate expression of cocaine CPP in male rodents^6,7^; however, it is not known whether this is mediated through striatal enkephalin. Given reported sex differences in acquisition of cocaine CPP^10^, we also wondered whether there were sex differences in the sensitivity to opioid receptor antagonists. To determine this, we tested whether the non-selective opioid receptor antagonist naloxone attenuates expression of cocaine CPP equally between male and female D2-PenkKOs and controls (Fig. 2A). After establishing cocaine CPP (Post Test 2), a preference test was performed following naloxone treatment (Post Test 3). Relative to preference at Post Test 2, naloxone significantly decreased preference for females, but not males, and this was independent of genotype (Sex x Test: F_1, 73_ = 9.31, *p* < 0.01; female: t_75_ = 5.36, *p* < 0.0001; Fig. 2B,C). Further, percent time on the cocaine floor at Post Test 3 was significantly lower when comparing females with males (t_158_ = 3.45, *p* < 0.01). These results were corroborated by additional analysis of the “Time on Grid” data (Table 1), and together suggest that naloxone attenuates expression of cocaine preference selectively in females by blocking the action of opioid peptide other than striatal enkephalin.

### Striatal enkephalin facilitates maintenance of cocaine preference during extinction

We next tested whether striatal enkephalin facilitates cocaine preference by preventing extinction and whether this differs by sex. After establishing cocaine preference and testing under naloxone conditions, mice underwent 8 forced-extinction trials where saline was paired with their previously cocaine (CS+) and saline (CS-) paired floors (Fig. 3A). After four extinction trials, all mice, regardless of genotype, extinguished their conditioned cocaine preference compared to the percent time on the cocaine floor at Post Test 2 (Test: F_2.33, 61.35_ = 23.20, *p* < 0.0001; t_26_ = 4.79, *p* < 0.001). Cocaine preference further decreased after four additional extinction trials relative to Post Test 2 (t_28_ = 6.79, *p* < 0.0001) and EXT Test 1 (t_26_ = 3.258, *p* < 0.05; Fig. 3B,C). While females consistently showed lower CPP than males (Sex: F_1, 25_ = 5.38, *p* < 0.05), we suspected this was a carryover-effect of acute naloxone attenuating preference in females at Post Test 3. Since this may mask a potential genotype effect, we calculated a CPP change score making preference relative to Post Test 3. Using this metric, we found that male and female D2-PenkKOs, but not *Penk*^*f/f*^ controls, decreased their cocaine preference between 4 and 8 extinction trials (Genotype x Test: F_1, 25_ = 8.067, *p* < 0.01; t_25_ = 4.192, *p* < 0.001; Fig. 3D). These data indicate that striatal enkephalin supports cocaine preference during extinction learning and may slow or prevent the extinction of cocaine-cue associations.

**Figure 3.**
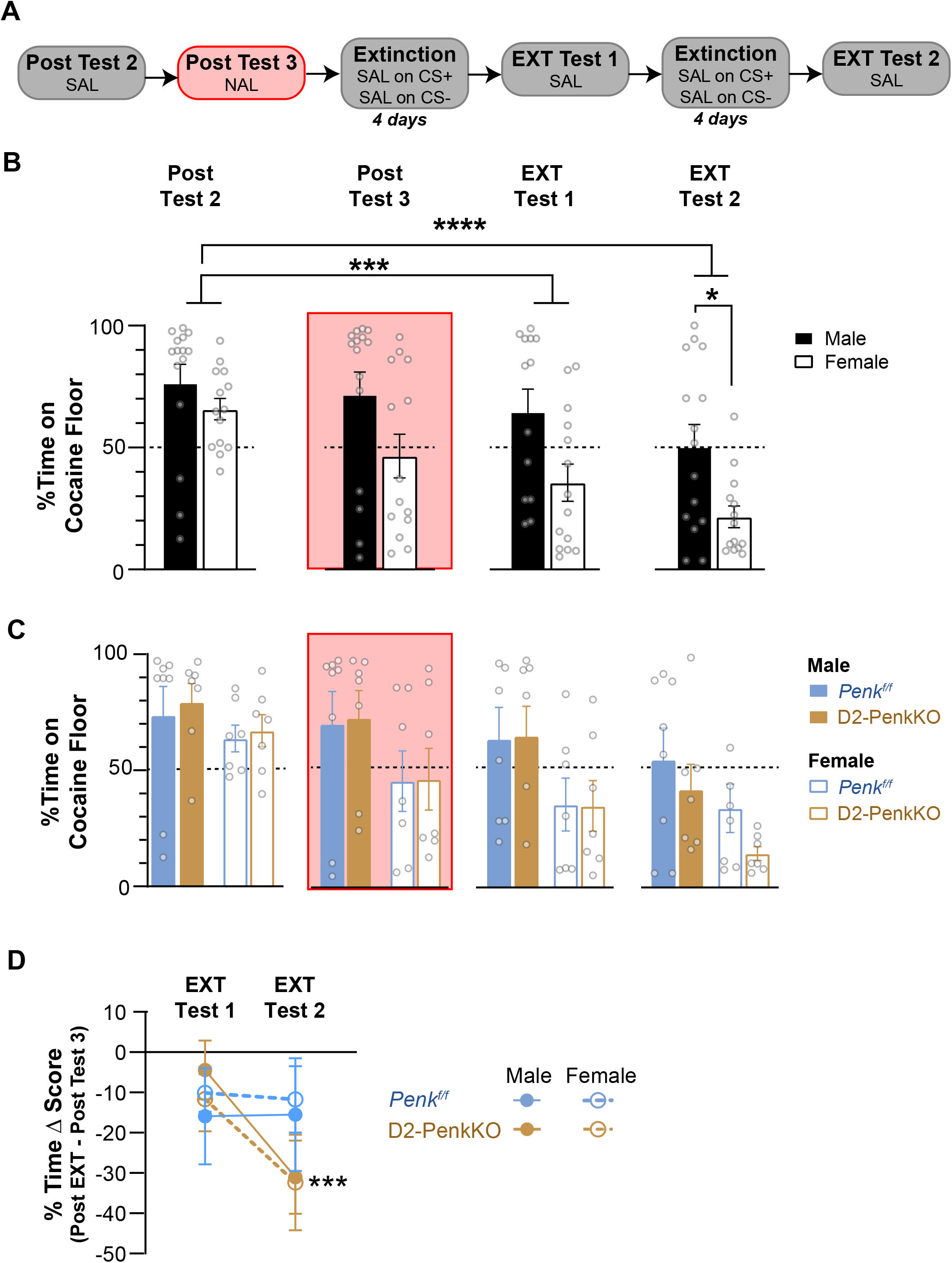
Striatal enkephalin facilitates maintenance of cocaine CPP. (A) Experimental timeline of cocaine CPP extinction training. All mice previously underwent a Pretest, 8 days of cocaine conditioning, and 3 preference tests (see Fig. 2A) prior to beginning extinction. *Penk*^*f/f*^ and D2-PenkKO mice received saline (SAL) 20 min prior to placement on the cocaine-paired floor (conditioned stimulus positive, CS+) during extinction training. Mice received 2 preference tests (EXT Test 1 and EXT Test 2) after 4 and 8 days of extinction. (B) Percent time on the cocaine floor decreased after 4 (*** *p* < 0.001) and 8 (**** *p* < 0.0001) extinction trials for both males (n = 15) and females (n = 14), regardless of genotype, relative to Post Test 2. (C) Percent time on the cocaine floor shown for each genotype (*Penk*^*f/f*^ and D2-PenkKO) and sex during Post Test 2, Post Test 3, EXT Test 1, and EXT Test 2 (n = 7 – 8 / genotype). (D) Change in the percent time on the cocaine floor (Post Ext test – Post Test 3) shows that male and female D2-PenkKO spend significantly less time on the cocaine floor during EXT Test 2 compared to EXT Test 1 (*** *p* < 0.001).

### Naloxone maintains cocaine preference during extinction in mice lacking striatal enkephalin

Systemic naloxone facilitates extinction of ethanol CPP^18^, and our data indicate that striatal enkephalin supports cocaine CPP during extinction. Thus, we hypothesized that naloxone facilitates extinction of cocaine CPP by blocking striatal enkephalin. To test this, *Penk*^*f/f*^ and D2-PenkKO mice were divided into two groups that received either saline (SAL on CS+) or naloxone (NAL on CS+) on their previously cocaine paired floor during forced-extinction trials (Fig. 4A).

**Figure 4.**
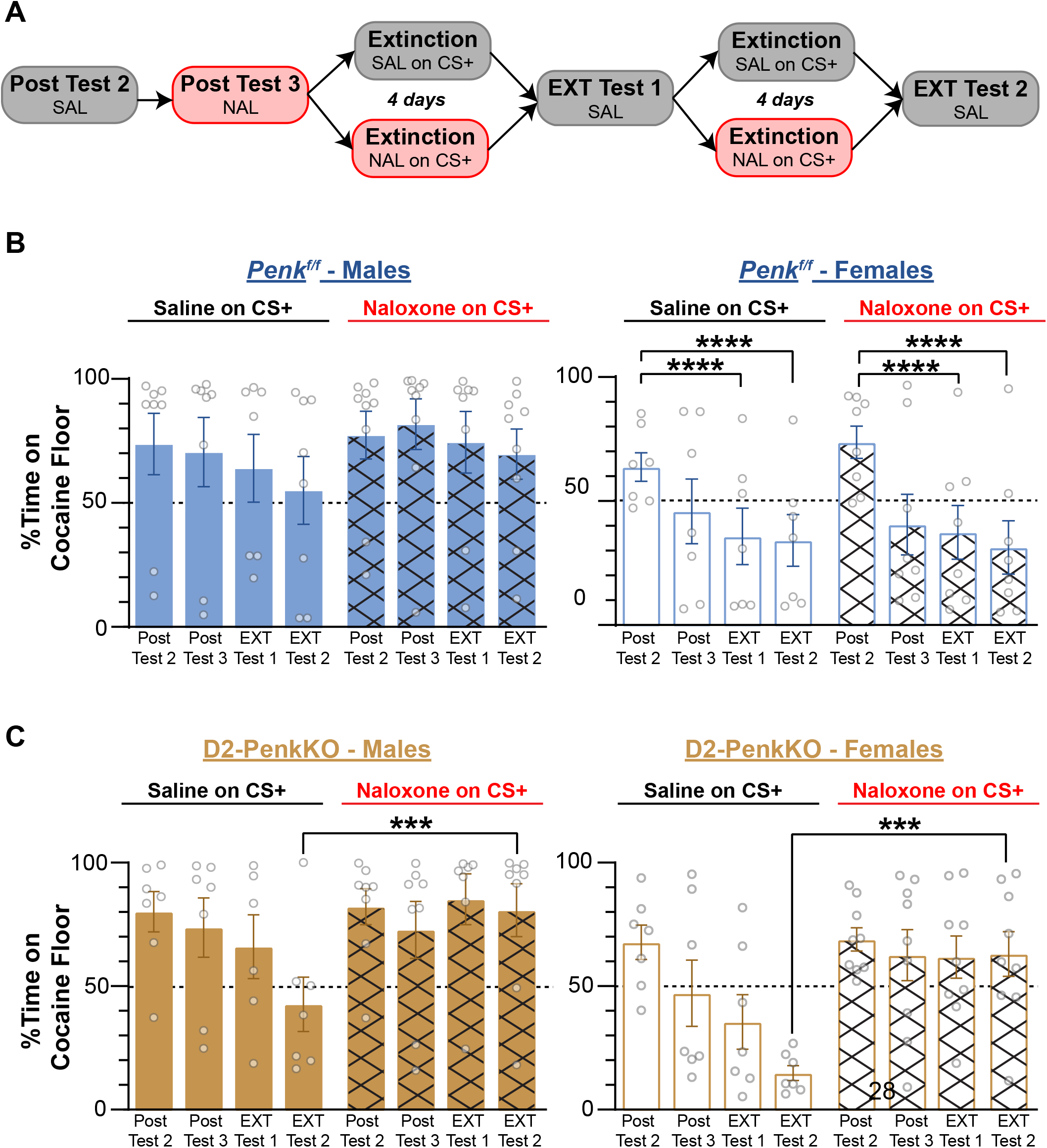
Naloxone maintains cocaine preference in mice lacking striatal enkephalin. (A) Timeline of cocaine CPP extinction training. *Penk*^*f/f*^ and D2-PenkKO mice either received naloxone (10 mg/kg; NAL) or saline (SAL) 20 min prior to placement on the cocaine-paired floor (conditioned stimulus positive, CS+) during extinction training. All mice received saline injection on the saline-paired floor (CS-) and 2 preference tests (EXT Test 1 and EXT Test 2) after 4 and 8 days of extinction. (B, left) Percent time on the cocaine floor was not reduced after 4 or 8 extinction trials for neither the saline-treated nor naloxone-treated male *Penk*^*f/f*^ mice (n = 8 saline, 9 naloxone). (B, right) Percent time on the cocaine floor was decreased after 4 or 8 extinction trials relative to Post Test 2 (\*\*\*\**p* < 0.0001) for both the saline-treated and naloxone-treated female *Penk*^*f/f*^ mice (n = 7 saline, 8 naloxone). (C) Naloxone-treated male (left) and female (right) D2-PenkKO mice maintained their cocaine preference after 4 and 8 extinction trials. Naloxone-treated male and female D2-PenkKOs had higher preference at EXT Test 2 compared to saline-treated D2-PenkKOs (\*\*\**p* < 0.001).

In control *Penk*^*f/f*^ mice, naloxone did not facilitate extinction for either sex compared to the saline group (Fig 4B). Both saline- and naloxone-treated female *Penk*^*f/f*^ mice showed extinction of preference (Sex x Test: F_3, 82_ = 4.94, *p* < 0.01) over four (t_14_ = 4.75, *p* < 0.001) and eight extinction trials (t_14_ = 5.25, *p* < 0.001) relative to pre-extinction preference levels at Post Test 2. In contrast, neither saline- nor naloxone-treated male *Penk*^*f/f*^ mice extinguished their preference, and male mice exhibited persistently stronger preference relative to females (EXT Test 1: t_27.23_ = 2.81, *p* < 0.05; EXT Test 2: t_29.86_ = 2.74, *p* < 0.05). Additionally, similar to the lack of effect of acute naloxone, repeated naloxone during extinction did not suppress preference in male *Penk*^*f/f*^ mice when analyzing the preference change score.

We next assessed how naloxone affected extinction of CPP in mice lacking striatal enkephalin. Surprisingly, naloxone increased preference in male and female D2-PenkKOs relative to the saline group after eight extinction trials (Naloxone x Test: F_3, 79_ = 11.13, *p* < 0.0001; t_114_ = 4.23, *p* < 0.001; Fig. 4C). Indeed, the degree of preference after extinction was similar to that before extinction (Post Test 2) for male and female D2-PenkKO mice. Interestingly, this preference-preserving effect of repeated naloxone was not seen in *Penk*^*f/f*^ controls, and preference was higher in naloxone-treated D2-PenkKO females than naloxone-treated *Penk*^*f/f*^ females (Genotype x Naloxone x Test, F_3, 81_ = 3.67, p < 0.05; t_27_ = 2.75, *p* < 0.05; Fig. 4B,C). While naloxone-treated male D2-PenkKOs had similar preference levels compared to naloxonetreated *Penk*^*f/f*^ males, we suspect this was because the naloxone-treated *Penk*^*f/f*^ males never extinguished. Indeed, analysis of the preference change score confirmed that naloxone treatment during extinction facilitates preference in both male and female D2-PenkKOs (Genotype, F_1, 28_ = 2.947, *p* < 0.05; S3).

## Discussion

In the present study, we generated a novel, cell-specific enkephalin knockout mouse to determine the role of striatal enkephalin in development, maintenance, and extinction of conditioned cocaine reward. Our findings suggest that although striatal enkephalin is not necessary for acquisition and expression of conditioned cocaine reward, it maintains cocaine preference, as D2-PenkKO mice show faster extinction of CPP. This is consistent with our previous work, which shows that enhanced met-enkephalin tone in the striatum potentiates acquisition of cocaine place preference^1^. Thus, striatal enkephalin appears to augment cocaine reward, and heightened striatal enkephalin tone may facilitate cocaine-cue associations and prevent extinction learning.

Non-selective opioid antagonists as well as MOR-selective antagonists attenuate expression of cocaine preference^2,6^. However, whether there are differences in the sensitivity to opioid antagonists between sexes remained unknown. Our experimental design enabled analysis of sex as a biological variable and revealed that naloxone attenuates expression of cocaine preference selectively in females. Multiple clinical trials have demonstrated mixed results in the efficacy of opioid medications for treatment of cocaine use^13,19–21^. It is possible these mixed results in part stemmed from sex differences in the response to opioid medications. Our results emphasize the consideration of biological sex in addiction treatment and suggest a single naloxone administration has long-lasting cocaine reward suppressing effects in females.

Single-dose naloxone also attenuated cocaine preference in D2-PenkKOs, suggesting that naloxone suppresses cocaine preference by blocking the action of a striatal opioid peptide other than enkephalin or that it is acting in a different brain region. The opioid peptide β-endorphin is released into the striatum, has high affinity for the MOR, and mice lacking β-endorphin show attenuated cocaine preference^22,23^. Systemic naloxone could also prevent enkephalin-mediated disinhibition of dopamine neurons in the ventral tegmental area^24,25^, which could potentially lead to attenuated cocaine preference^2^.

Despite naloxone’s acute reward-suppressing effects in females, repeated administration during extinction trials did not facilitate extinction of cocaine preference in *Penk*^*f/f*^ mice. In females, we suspect a naloxone-effect was masked by a floor effect. However, in males this finding is consistent with previous research that shows male DBA/2J are resistant to naloxone-induced facilitation of cocaine CPP extinction^18^. Together, these data suggest wildtype females are uniquely sensitive to the reward-blocking effect of naloxone, while males are robustly resistant.

Our findings also indicate that striatal enkephalin is important for maintaining the cocaine-context association during extinction. Mice lacking striatal enkephalin extinguished cocaine preference faster and to a greater degree than littermate controls. This is consistent with previous research showing that enhanced striatal enkephalin tone facilitates acquisition of a cocaine-context association^1^. Quite surprisingly, though, naloxone treatment during extinction trials facilitated preference and prevented extinction in mice lacking striatal enkephalin. Thus, it appears that naloxone prevented extinction by blocking a different opioid peptide. One potential candidate is dynorphin, opioid peptide associated with stress and the negative affective state related to cocaine withdrawal^26^. Deletion of striatal *Penk* may disrupt the balance between enkephalin and dynorphin within the striatum, resulting in a relatively higher dynorphin tone and a shift away from cocaine reward and toward extinction of preference. Thus, naloxone treatment during extinction in D2-PenkKO mice would be expected to block the predominant dynorphin signal and bias towards maintaining cocaine preference. This hypothesized mechanism is consistent with reports showing mice lacking *Pdyn* have slower extinction of cocaine self-administration^4^. Although we did not detect heightened striatal *Pdyn* mRNA levels in D2-PenkKOs, future studies should evaluate whether these mice show altered dynorphin peptide levels or whether dynorphin contributes to this behavioral effect.

Taken with the previous literature, these data indicate a critical role for enkephalin in maintaining the drug-context associations that drive cocaine place preference. While not necessary for acquiring these associations, enkephalin appears to augment the rewarding effect of cocaine to sustain cocaine seeking in a Pavlovian paradigm. Further, wildtype females are uniquely susceptible, while males are resistant, to the reward suppressing effects of acute naloxone treatment, which could have treatment implications for clinical populations. Conversely, repeated treatment with naloxone in an extinction paradigm actually prevents extinction in subjects with low striatal enkephalin. This may be an important consideration for treatment of clinical populations because we previously demonstrated that withdrawal from short-term cocaine downregulates striatal *Penk*^1^. Thus, significant reduction of striatal enkephalin may occur in human subjects with casual, recreational use of cocaine as well as in subjects with single nucleotide polymorphisms in the *Penk* loci. Based on our findings, repeated treatment with opioid receptor antagonists, particularly naltrexone, may be contraindicated for those being treated for alcohol or opioid use disorder with concurrent cocaine use.

## Data Availability Statement

The data that support the findings of this study are available on request from the corresponding author.

## Acknowledgments

This research was supported by a grant from the National Institute of Drug Abuse to LKD (R01DA054329), a Rising STARs Award from the University of Texas at Austin to LKD, and a Bruce Jones Predoctoral Fellowship to KM. We are grateful to Dr. Andreas Zimmer for generously providing the floxed *Penk* mice and to Dr. Ian Riddington and the Mass Spectrometry Facility at UT Austin for scientific assistance.

## Author Contributions

KM designed and performed most of the experiments, conducted data analysis, and prepared the manuscript. IBC assisted in RNAscope and qPCR experiments. MA, NNL, and LN performed experiment and data analysis for mass spectrometry. We very much appreciate AZ for graciously providing *Penk*^*f/f*^ strain with us. LKD assisted in experimental design, data analysis, and manuscript preparation.

## Competing Interests

We have no competing interests to report.

## Notes

### Competing Interest Statement

The authors have declared no competing interest.

