## Supplemental Figures for "Striatal enkephalin supports maintenance of conditioned cocaine reward during extinction"

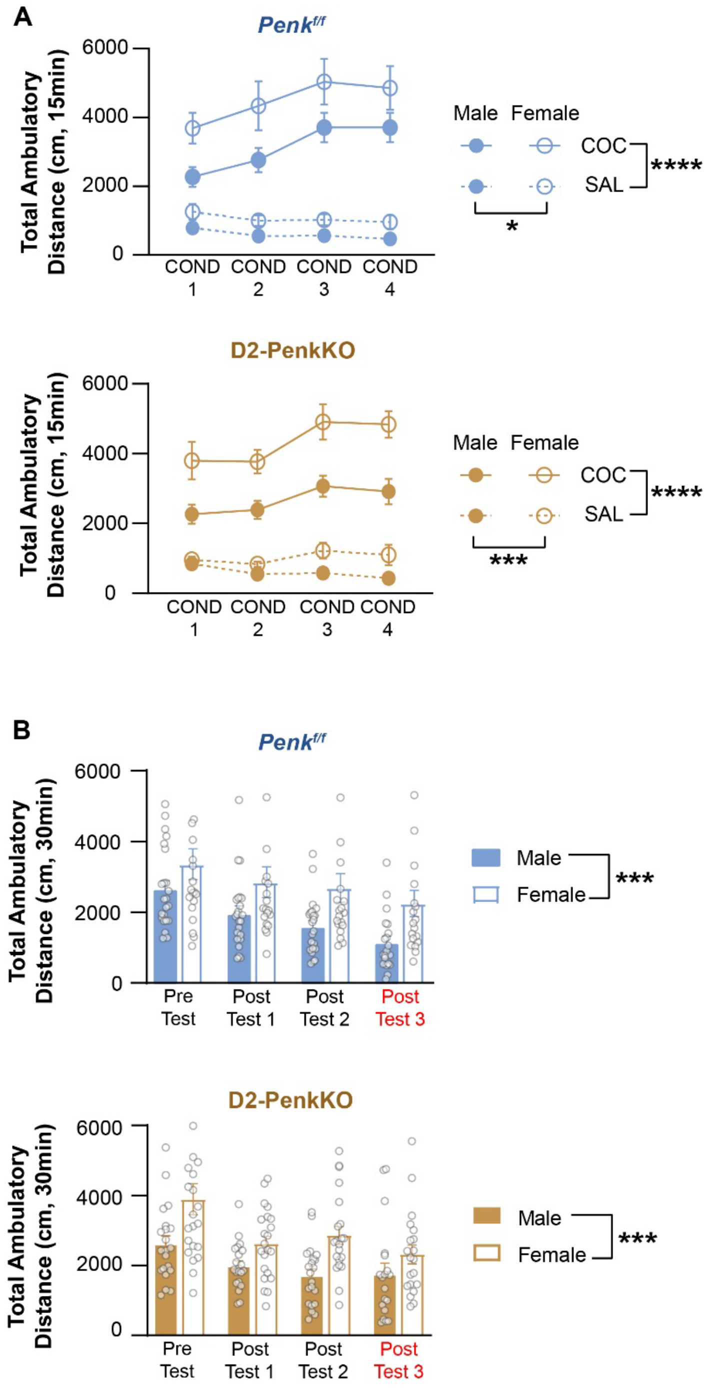


**Supplemental Figure 1. Locomotor activity during cocaine place conditioning and testing.** (A) Total distance *Penk$^{f/f}$* and D2-PenkKO traveled during 15 min conditioning sessions. All mice significantly traveled less during the 3$^{rd}$ and 4$^{th}$ saline sessions compared to the 1$^{st}$ session. Cocaine increased locomotor activity on sessions 3 and 4 relative to the 1$^{st}$ session for all mice. Females consistently showed greater activity than males, regardless of genotype. (B) Total distance *Penk$^{f/f}$* and D2-PenkKO traveled during 30 min preference tests. All mice decreased activity during Post Tests 1, 2 and 3 relative to Pre-Test. Females were more active than males across all tests, regardless of genotype.

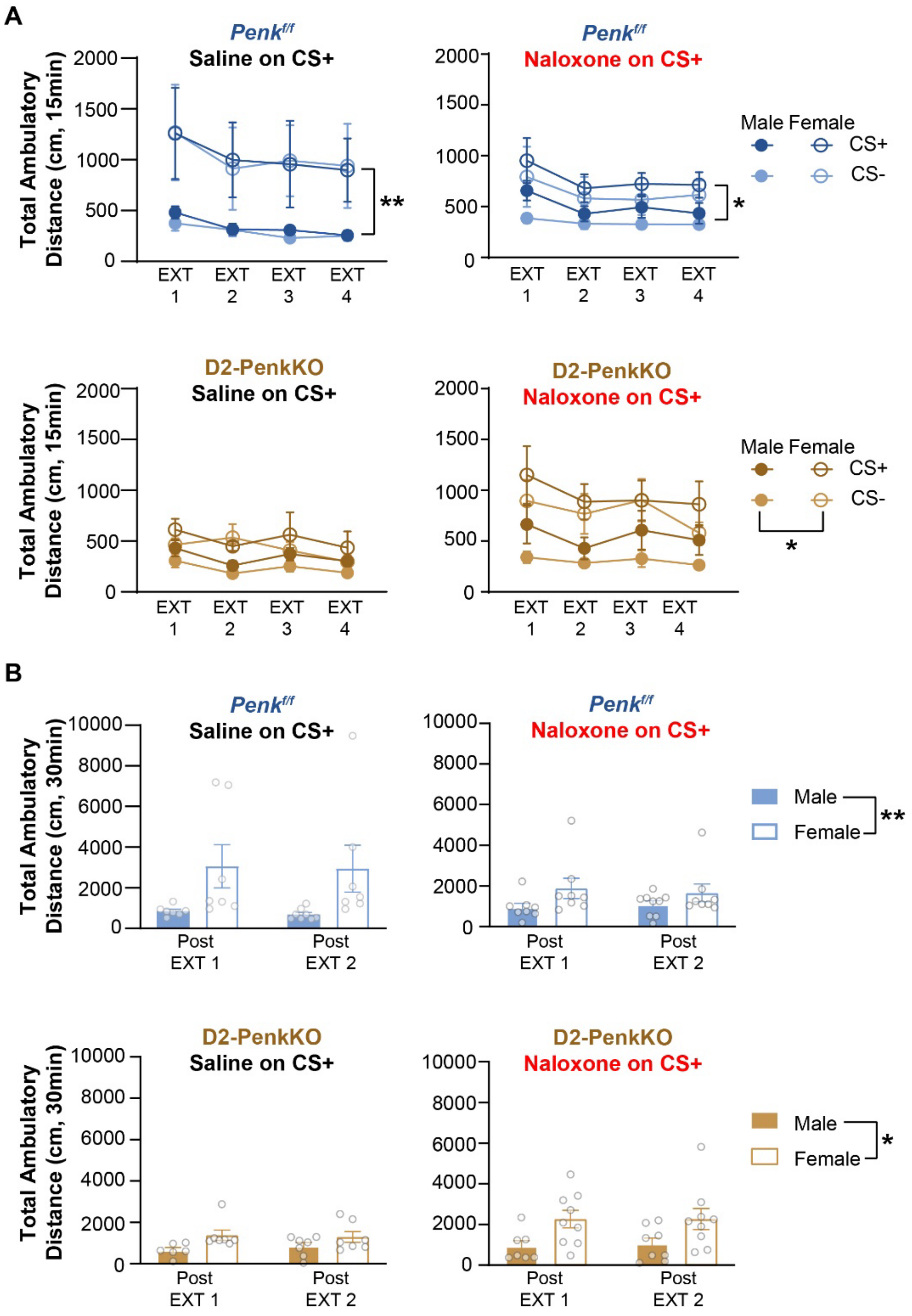


**Supplemental Figure 2. Locomotor activity during extinction trials and post-extinction preference tests.** (A) Total distance *Penk*$^{f/f}$ (top) and D2-PenkKO (bottom) traveled during 15 min extinction trials. All mice significantly decreased activity over the course of 2 weeks of extinction training. There were no differences between genotype or pretreatment group (saline or naloxone). Females were consistently more active than males, regardless of genotype. (B) Total distance *Penk*$^{f/f}$ (top) and D2-PenkKO (bottom) traveled during 30 min preference tests. There were no differences between genotype or pretreatment group (saline or naloxone). Females were consistently more active than males, regardless of genotype.

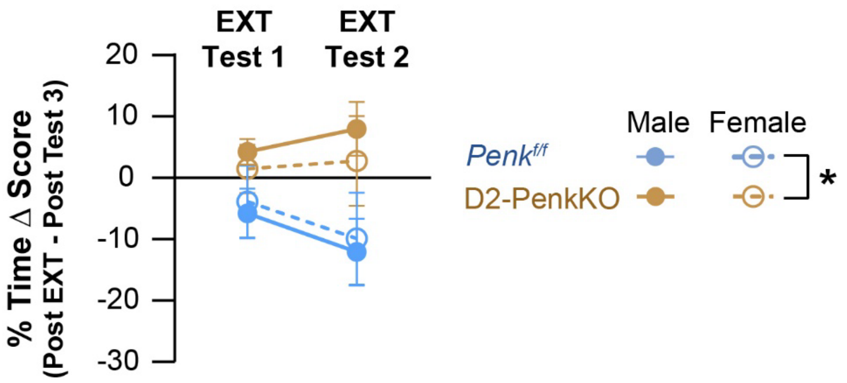


**Supplemental Figure 3. Change in percent time on the cocaine paired floor following naloxone-paired extinction.** The change in percent time spent on the cocaine floor is shown for Extinction Tests 1 and 2. *Penk*$^{f/f}$ and D2-PenkKO mice received naloxone paired with the previous cocaine-paired floor during extinction trials over 8 days. Both male and female D2-PenkKO mice spent more time on the previous cocaine-paired floor relative to *Penk*$^{f/f}$ littermate controls at both extinction tests. There was no effect of sex for either genotype.
